# Memory recall errors reflect interacting sensory and mnemonic representations

**DOI:** 10.1101/2025.10.20.683560

**Authors:** Holly Kular, John T. Serences

## Abstract

Visual working memory (WM) enables the maintenance of information that is no longer present in the environment. Some accounts propose that WM is supported by abstract representations so that new sensory inputs do not interfere with existing memories. Others posit that early sensory representations are recruited to maintain memory precision, potentially at the cost of interference caused by new inputs. Here we tested these accounts using an orientation recall task to determine whether memory errors reflect interacting representations of sensory and mnemonic information. We found that adding noise to the memoranda and presenting feature-neutral distractors independently increased recall errors. However, distractors that shared a feature with remembered stimuli led to systematic attractive biases such that memory errors were pulled toward the orientation of the distractor. The magnitude of this bias was modulated by both stimulus noise and whether the distractor was behaviorally relevant. Our results demonstrate that while working memory can utilize abstract representations, it remains susceptible to feature-specific sensory interference, suggesting partial reliance on sensory-like codes.

## Introduction

Visual working memory flexibly supports a wide range of cognitive tasks by maintaining information that is no longer present in the environment to guide decision making and action planning. There are two prominent theoretical frameworks regarding how visual working memory (WM) representations are maintained in the brain. One perspective argues that WM is supported primarily by neural populations in frontal and parietal cortices that maintain representations that are abstracted away from low-level sensory features (Bettencourt & Xu, 2016; Fuster & Alexander, 1971; Goldman-Rakic, 1995; Mendoza-Halliday et al., 2014; Riley & Constantinidis, 2015; Xu, 2017, 2020, 2024). An alternative, and not mutually exclusive, view suggests that WM recruits representations in the same regions of early visual cortex that are involved in early perceptual processing (D’Esposito, 2007; D’Esposito & Postle, 2015; Harrison & Tong, 2009; Serences, 2016; Serences et al., 2009). That said, frameworks that allow for sensory recruitment and those that do not make distinct predictions about how memory representations interact with new sensory inputs. If WM relies exclusively on abstract formats in frontal-parietal regions, then ongoing sensory processing should not interfere with stored mnemonic information (Bettencourt & Xu, 2016; Mendoza-Halliday et al., 2014; Xu, 2017, 2020, 2024). However, if early visual cortex plays an active role in WM, then sensory and mnemonic representations may share neural resources, potentially leading to systematic interactions between stored memories and new inputs.

Previous fMRI research has shown that mnemonic information is encoded in activation patterns in visual cortex, demonstrating that memory representations are maintained in early visual cortex even after the stimulus input has ended (Christophel et al., 2018; Harrison & Tong, 2009; Rademaker et al., 2019; Serences, 2016; Serences et al., 2009; Sprague et al., 2014, 2016; Sreenivasan et al., 2014). While the structure of mnemonic and sensory representations may not be identical (Libby & Buschman, 2021), early sensory areas may play an important role when remembering fine details, as neurons in these areas are capable of supporting high-precision representations of relevant information (Harrison & Tong, 2009; Serences, 2016; Serences et al., 2009). Moreover, other studies have shown that distractors can interfere with and attractively bias representations of remembered stimuli, particularly when distractors and the memoranda come from the same feature domain (Dubé et al., 2014; Hallenbeck et al., 2021; Huang & Sekuler, 2010; Lorenc et al., 2018; Mallett et al., 2020; Nemes et al., 2011, 2012; Rademaker et al., 2015; Teng & Kravitz, 2019; Van der Stigchel et al., 2007; Zhang & Lewis-Peacock, 2023). These behavioral effects may arise from interacting, or *entangled,* representations of sensory and mnemonic information (DiCarlo et al., 2012; DiCarlo & Cox, 2007; Pagan et al., 2013). Consistent with this account, Hallenbeck et al. (2021) observed that behavioral reports of a remembered spatial position were attracted to a nearby distractor, as were neural representations of the remembered position in early visual areas V1–V3 as assessed with fMRI.

However, given that distractor resistance is a key feature of WM (Lorenc et al., 2021), some have reasonably argued against sensory recruitment on the basis that repurposing sensory neurons to encode new inputs and mnemonic information could lead to interference and confusion between seen and remembered stimuli (Bettencourt & Xu, 2016; Mendoza-Halliday et al., 2014; Xu, 2017, 2020). This is an important concern because we typically move our eyes several times a second and new sensory inputs are continuously being processed and may overwrite mnemonic representations in early visual cortex. Indeed, prior studies using complex objects have reported little effect of irrelevant distractors on memory recall error or on neural representations in early visual cortex (Bettencourt & Xu, 2016; Xu, 2024). Some propose that WM is supported by more abstract codes in parietal cortex that are reformatted so that they are ‘untangled’ from the feature-specific content of new sensory inputs (Xu, 2024).

Here we test whether sensory-like codes can play a role in supporting WM by determining whether sensory and mnemonic representations systematically interact. In a simple recall task for a remembered orientation, we presented either no distractor or a distractor that could be behaviorally relevant. Additionally, we manipulated the sensory uncertainty of the memory sample as a means of parametrically varying memory strength. By using a well-characterized stimulus space we could make precise predictions about the form of potential interactions between sensory and mnemonic representations (e.g. attractive biases between remembered and seen orientations). Moreover, by varying sensory uncertainty we could evaluate the relationship between memory fidelity and susceptibility to bias and we could establish sensitivity to detect distractor-induced changes in recall error should any exist. For example, prior work demonstrates that changing the amount of coherent information in a stimulus reduces the precision of feature-selective population codes in early visual cortex (Newsome et al., 1989; Parker & Newsome, 1998) and that attending to specific features, such as the orientation of a distractor, increases the gain of neurons in early visual cortex that are tuned to the relevant feature (McAdams & Maunsell, 1999; Moran & Desimone, 1985; Reynolds & Chelazzi, 2004; Reynolds & Heeger, 2009; Treue & Martínez Trujillo, 1999). Thus, manipulating sensory noise and distractor relevance should lead to co-occurring modulations in early visual cortex, affording the opportunity to test for interactions between seen and remembered stimuli. If mnemonic representations are abstracted and protected from distractor interference via untangling, then we predict larger recall errors due to changes in uncertainty but no systematic interactions between memory and sensory representations. However, if WM is dependent on sensory-like representations, then we predict that errors should be systematically biased by the presence of feature-similar distractors, particularly when memory strength is low and distractors are behaviorally relevant.

## Experiment 1

We first designed an experiment to assess whether increased sensory uncertainty associated with a memory stimulus leads to more susceptibility to interference from distractors presented during the delay period. We collected data from 42 participants (21 female, mean age = 20.6), from the University of California San Diego who completed the study for course credit or monetary award ($15/hour). The protocol for this study was approved by the Institutional Review Board at UCSD, and all participants provided written informed consent. Participants were excluded if they failed to respond on more than 10% of all trials. After excluding 8 subjects, all analyses were conducted on the remaining 34.

### Task

On each trial, a phase-reversing oriented ‘sample’ stimulus, rendered in a circular shape (7° radius) was presented for 500 ms, with a circular annulus around a central fixation point cropped out of the stimulus (2° radius) (Figure 1A). The orientation information within the stimulus was parametrically manipulated with a bandpass filter with the width determining the magnitude of orientation information in the stimulus. First, we generated a white-noise image that was 520 x 520 pixels by making random draws from a normal distribution with a mean of zero and a variance of 1. The image was then transformed into the frequency domain (using *fft2.m*). We then multiplied the frequency-domain representation with an orientation filter consisting of a circular Gaussian (Von Mises) function centered at the desired orientation, with concentration parameters (k) of 50, 100 and 5000 for the low, medium and high-noise stimuli, respectively (Figure 1B). We then applied a spatial frequency bandpass filter from 1 to 4 cycles/pixels, with Gaussian smoothed edges (smoothing SD = 0.005 cycles/pixel). After multiplying the frequency domain representation of the white noise stimulus with these filters, we replaced the image’s phase information with random values uniformly sampled between –pi to +pi before transforming back into the spatial domain (using *ifft2.m*). Finally, we multiplied the image by a scaling factor of the noise image contrast divided by the filtered image contrast in order to account for some of the contrast lost during filtering.

**Figure 1.**
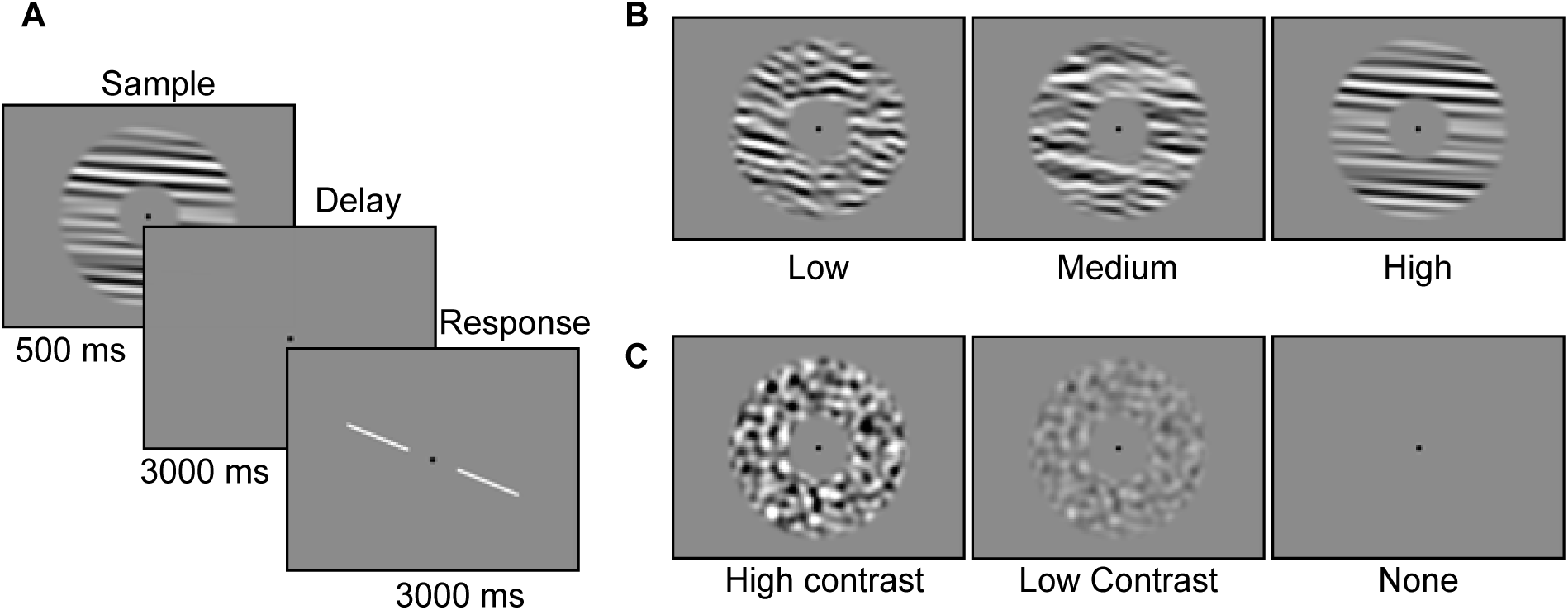
(A) Trial structure. Example displays for stimulus noise (B) and distractors (C).

Participants encoded the orientation of each sample stimulus and retained the information across a 3000 ms delay period. During the delay, participants saw either a blank screen (no distractor), a 50% Michelson contrast filtered noise distractor, or a 100% Michelson contrast filtered noise distractor (Figure 1C) (where the distractors had no bias in orientation information but were filtered to have the same mean spatial frequency composition as the memory sample stimulus following the procedure described above). After the retention interval, subjects rotated a white probe line using four buttons on a standard keyboard to match the remembered orientation of the sample stimulus.

### Procedure

The sample stimulus orientation was first chosen from one of six, 30° orientation bins that uniformly spanned the 180° stimulus space. Sample stimuli were drawn from each of the 6 bins equally often during a run and once the bin was determined on a trial, the exact orientation was drawn from a uniform distribution across all angles in that bin to ensure that the stimuli evenly tiled orientation space. The orientation of the response probe was chosen randomly with respect to the orientation of the sample stimulus on each trial. Each run consisted of 36 trials and there were 10 runs for a total of 360 trials per participant and stimulus noise, orientation, and distractor type were fully counterbalanced across runs. Participants were instructed to fixate on the central point throughout each trial and uniformly sampling across the entire 180° orientation space discouraged verbal encoding strategies.

### Results

Working memory errors were quantified using the root mean squared error (RMSE) of the reported orientation compared to the actual orientation of the memory sample (Figure 2A). Bayesian repeated measures ANOVA revealed larger errors with higher stimulus noise (BF_10_ = 4.67x10^15^ ± 2.73%, η_p_^2^ = 0.27) and larger errors associated when distractors were present (BF_10_ = 27.8 ± 2.83%, η_p_^2^ = 0.05). Stimulus noise and distractor type did not interact to increase memory errors (BF_10_ = 0.05 ±2.82%, η_p_^2^ = 0.01). A post-hoc Bayesian t-test supports the null hypothesis that overall RMSE did not increase as a function of increasing distractor contrast (BF_10_ = 0.18 ±- 0.1%, Cohen’s d = 0.1).

**Figure 2.**
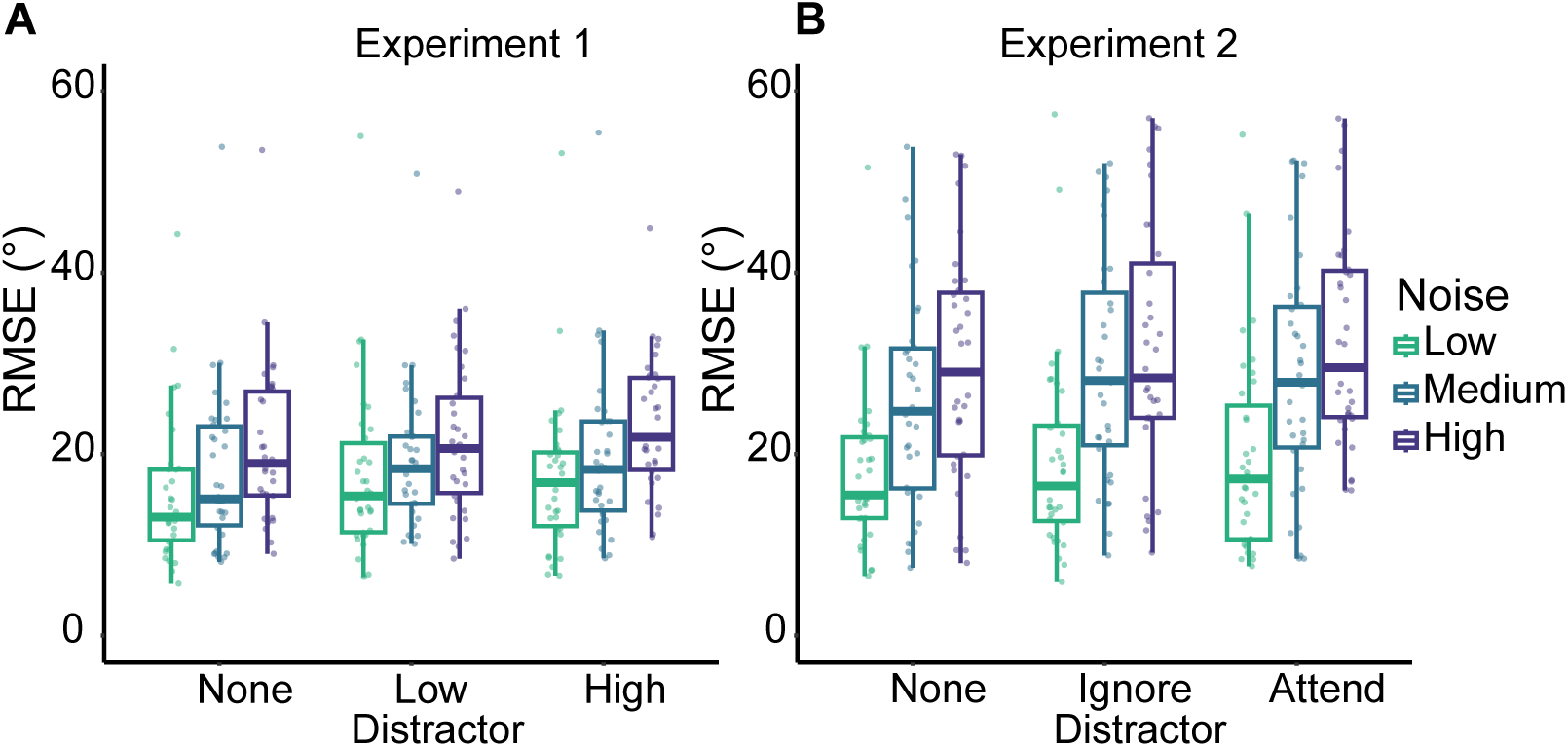
(A-B) Boxplots of response errors (RMSE). Plots are in the style of Tukey, boxes display interquartile range and whiskers are 1.5 x quartiles. Individual dots represent individual participant errors.

## Experiment 2

The second experiment assessed whether sensory uncertainty interacts with noise distractors when the distractors are behaviorally relevant. We collected data from 43 participants (33 female, mean age = 20.0) from the University of California San Diego who completed the study for course credit or monetary award ($15/hr). The protocol for this study was approved by the Institutional Review Board at UCSD, and all participants provided written informed consent. Participants were excluded if they failed to respond on more than 10% of all trials. After excluding 8 subjects, all analyses were conducted on the remaining 35.

### Task & Procedure

Experiment 2 followed the same task and procedure as Experiment 1 except for the following modifications to the distractor. The distractor condition was either none, ignore, or attend. On ignore and attend trials, the distractor was always 65% Michelson contrast noise mask. On the ignore-distractor trials, subjects passively viewed the distractor, as in Experiment 1. On the attend-distractor trials, subjects were instructed to press a key in response to a brief decrease in the contrast of the distractor that occurred for 125 ms at a uniformly and pseudo-randomly distributed time between 500 ms and 2500 ms of the 3000 ms delay period.

### Results

As in Experiment 1, response errors (Figure 2B) were larger with more stimulus noise (BF_10_ = 4.60 x 10^33^ ±1.45%, η_p_^2^ = 0.45), and distractor presence (BF_10_ = 34.9 ± 1.86%, η_p_^2^ = 0.07). Although, a post-hoc Bayesian t-test did not support that the addition of having to attend and respond to the distractor increased error (BF_10_ = 0.11 ± 0.15%, Cohen’s d = 0.005). Finally, the Bayesian repeated measures ANOVA supports the null hypothesis that stimulus noise and distractors do not interact to magnify memory errors (BF_10_ = 0.03 ± 1.73%, η_p_^2^ = 0.001). Even though we repeatedly observed a null interaction between stimulus noise and distractor interference in the magnitude of WM errors, this may reflect the operation of an unobserved non-linearity that compresses the combined impact of sensory noise and distractor interference on behavioral measurement like RMSE. For example, there are reducible and irreducible interactions: the former can be undone by a transform of the data, whereas the latter are cross-over interactions and generally cannot be undone (Loftus, 1978). Therefore, the lack of an interaction might be due to a non-linearity that reduces a real interaction at the level of the generating mechanism into an effect on the measurement that appears to be additive.

## Experiments 3 and 4

Experiments 1 and 2 established that the presence of a feature-neutral distractor led to higher error rates overall, with no interaction between distractor salience/relevance and stimulus noise. The lack of an interaction is consistent with models suggesting that mnemonic and sensory representations are ‘untangled’ from sensory representations, perhaps via reformatting of the remembered information for storage in higher-order cortical areas (Bettencourt & Xu, 2016; Xu, 2024). However, systematic interactions between memory and sensory stimuli could still exist, such as mean-reverting attractive or repulsive biases, without inflating the RMSE and giving rise to an interaction between stimulus noise and distractor salience/relevance. Indeed, prior studies that have investigated interference have often used different categories of objects as stimuli (Xu, 2024), and our Experiments 1 and 2 used a filtered noise distractor that did not contain any coherent orientation information. Thus, many prior studies – including the present Experiments 1 and 2 – were not optimized to assess interactions between memory and sensory stimuli that may reveal entangled representations even in the absence of effects on summary measures like RMSE.

To provide a more direct and sensitive measure of interactions, Experiment 3 assessed whether sensory uncertainty interacts with noise distractors that are both behaviorally relevant and from the same feature domain as the remembered stimulus. Experiment 4 was a replication of Experiment 3 and both were pre-registered (Experiment 3: https://osf.io/k76dh/overview?view_only=0fa0f580d7a443629c21c60cd780b763; Experiment 4: https://osf.io/eafkd/overview?view_only=c0b92b8534084773bae496c0fed21189). Experiment 3 had 24 participants (15 female, mean age = 22.1) who completed 576 trials each. Experiment 4 had 12 participants (7 female, mean age = 23.4) who completed 1600 trials each. We collected data from participants at the University of California San Diego who completed the study for course credit or monetary award ($15/hr). The protocol for this study was approved by the Institutional Review Board at UCSD, and all participants provided written informed consent. We initially did not conduct a parametric power analysis prior to pre-registering Experiment 4 because our planned analyses were non-parametric. For Experiment 4, we initially estimated that with greater than double the number of trials per subject used in Experiment 3, we would be able to detect effects the size of Experiment 3’s with half the participants and so we pre-registered 12 subjects. However, after conducting a post-hoc parametric power analysis using the actual data from Experiment 3, we determined that the goal sample size should have been 17 participants. After reporting the results from Experiment 4 with the pre-registered sample size of 12, we report results after adding 5 more participants to Experiment 4a dataset for a total of 17 participants (11 female, mean age = 26.2). To clearly delineate the pre-registered version of Experiment 4 from the version with the post-hoc addition of 5 more subjects, we label the two versions Experiment 4a and 4b respectively.

### Task & Procedure

Experiments 3 and 4 were similar to Experiments 1 and 2 but with a different type of distractor and a modified task to perform on the distractor. The distractor was now an oriented rectangle that was 3.5° wide and 14° long that contained the same dynamic contrast filtered white noise as described in the previous experiments (Figure 3A). The rectangle orientation and sample stimulus orientation were independently selected from one of six, 30° orientation bins that uniformly spanned the 180° stimulus space and this was counterbalanced across runs. On each trial the distractor orientation briefly (125 ms) shifted either clockwise or counterclockwise. The size of the shift was staircased across runs and initialized at 10° on the first run and decreased by 1° down to a floor of 2° if distractor task accuracy was greater than 79% and increased by 1° if distractor task accuracy was less than 71% up to a ceiling of 20° (mean accuracy = 90.8% ± 7.96%). Experiment 3 had no-distractor, ignore-distractor, and attend-distractor conditions that were pre-cued at the start of each trial by changing the color of the fixation point to one of three colors (pre-cue presented for 1000 ms, and colors counterbalanced across participants). On attend-distractor trials, the distractor task was now a change discrimination task where the participants were asked to report whether the distractor rotated clockwise or counter-clockwise using the arrow keys on the keyboard. Since we had a well-established distractor effect in the previous experiments, we dropped the no-distractor condition for Experiment 4. Additionally, for Experiment 4 we sampled the distractor orientation with respect to the stimulus orientation in four 30° wide bins centered at 0°, -45°, +45°, and 90° away from the stimulus orientation. For Experiment 4, to ensure task comprehension, participants were required to correctly describe the instructions at the beginning of each data collection session, as well as given feedback during an observed practice block of trials on the first day of data collection.

**Figure 3.**
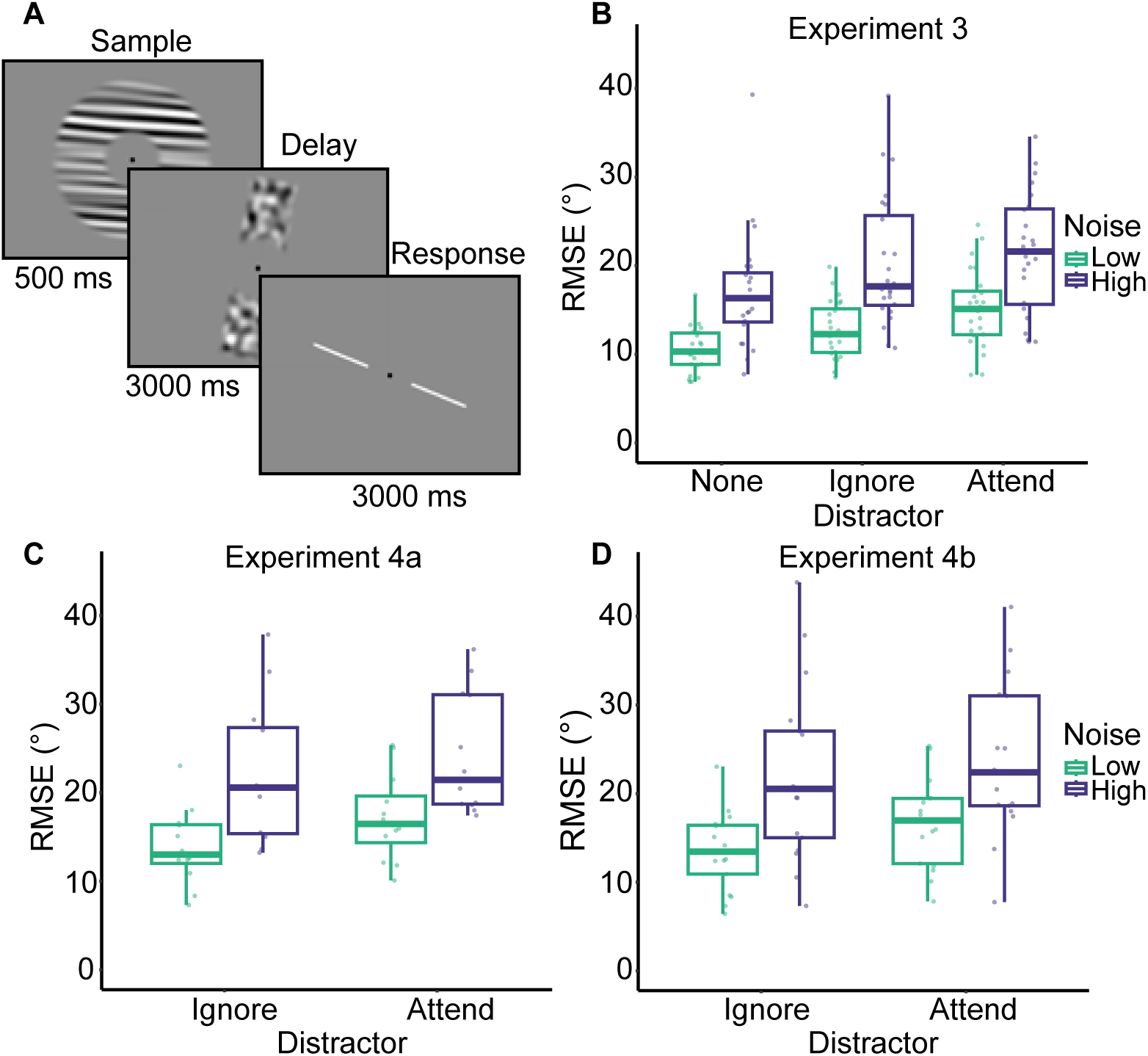
(A) Task structure for experiments 3-4. (B-D) Boxplots of response errors for Experiments 3 (B), 4a (C), and 4b (D).

### Pre-registered Exclusion Criteria

Participants were excluded if they failed to respond to the main orientation report task for 90% of all trials. Participants were also excluded if they failed to respond to the distractor task for 90% of all trials with a distractor task. Participants were excluded if they responded during the delay on more than 10% of the trials where there was no distractor task. Data collection continued until the pre-registered target sample size of 24 participants was reached (with 10 participants replaced for Experiment 3). For Experiment 4a, data collection continued until the pre-registered goal sample size of 12 participants was reached (with 3 participants replaced). For Experiment 4b, data collection continued for the additional 5 participants needed to reach the new goal sample size (with 4 participants replaced). Data collection was discontinued for participants if they exceeded the allowed number of failed responses for the whole experiment within the first day of data collection. During data collection, we noted the number of excluded participants and in hindsight the pre-registered exclusion criteria appeared to be too strict, but we opted to continue as planned to ensure participants were on-task. For Experiment 3, 9 participants were replaced for failure to respond correctly to the distractor task and 1 participant was replaced for failure to respond to the main task. For Experiment 4a, 3 participants were replaced for failure to respond correctly to the distractor task. For Experiment 4b, 4 participants were replaced for failure to respond correctly to the distractor task.

### Experiment 3: Fitting a Gaussian to folded response errors

Because we used a parametrically varying stimulus space and memory and distractor stimuli shared a common feature (orientation), we next evaluated bias in the memory response errors that was systematically linked to the oriented distractor. We quantified bias by computing the circular mean of the error distributions to infer the direction of the response errors with respect to the memory-distractor orientation offset on each trial. The memory-distractor orientation offset was defined as the circular mean difference in orientation between the distractor orientation and the sample stimulus, (𝛥𝜃 = 𝜃_D_ − 𝜃_S_). We performed this analysis on the continuous memory response errors from Experiment 3 on a moving window of 30°, such that a bias centered at 0° would include all trials with a 𝛥𝜃 in the range [-15°, 15°]. Before estimating bias, we first removed cardinal biases by fitting a 4th degree polynomial to the response errors arranged by sample orientation and we used the residuals from that fit for further analysis. We calculated both a forward (starting at -90°) and backward moving window (starting at 89°) and all analyses were conducted on the average of the two. We then folded the response errors by multiplying the trial-wise errors by the sign of 𝛥𝜃 and averaging. We fit a Gaussian parameterized with an amplitude 𝑎, width 𝑤, and center of the peak 𝑏 = 45°.

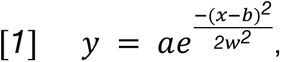

To fit our participant responses, x is 𝛥𝜃. We fit our Gaussian function using a bootstrap resampling method. First, we generated a sample of 24 pseudo-subjects by iteratively taking the average of a random sample of 24 subjects with replacement from the real subject data. Next, we calculated a repeated measures ANOVA on this sample of pseudo-subjects. We repeated this 5,000 times to construct a sample of ANOVA statistics. In order to create a null-distribution of ANOVA statistics for a permutation test, we scrambled the stimulus noise and distractor conditions in the following manner. To test the main effect of attention to the distractor, the null distribution was constructed with the distractor condition labels swapped. To test the main effect of stimulus noise, the null distribution was constructed with the stimulus noise labels swapped. To test the interaction between the stimulus noise and distractor attention, the null distribution was constructed with all labels randomly shuffled. For each sample fit, there were two free parameters: 𝑎 and 𝑤, which were optimized by minimizing the root mean squared error (RMSE) between the pseudo-subject average and the fit. We performed a grid search of our parameters and subsequently performed a local search around the most successful grid point using the Nelder-Mead algorithm up to 1x10^8^ iterations. Optimization of the parameters was bounded at (𝑎[-5, 8], 𝑤[20, 35]).

### Experiment 4: Response error bias estimation

Because we sampled distractor orientations centered on four distractor-sample (𝛥𝜃) bins (0°, - 45°, +45°, and 90°), we do not fit a continuous Gaussian function as we did for Experiment 3. Instead, mean response errors in each bin were folded such that the trial-wise errors in the -45°- centered bin were multiplied by -1 and averaged with the +45°-centered bin and the 0°- and 90°-centered errors were averaged. This results in average errors in two 𝛥𝜃 bins – oblique (-45° and +45°) and orthogonal (0°and 90°). To estimate amplitude of the response bias we subtracted mean response errors of the orthogonal bin from the oblique bin. We estimated the amplitude response bias using the same bootstrap resampling method as described in Experiment 3.

### Results of Experiments 3 and 4

#### Mean memory error

As shown in Figure 4B and 4D, working memory errors were larger with more stimulus noise (Figure 3B, Experiment 3: BF_10_= 8.92x10^14^ ± 3.15%, η_p_^2^ = 0.47; Figure 3C, Experiment 4a: 4.69x10^3^ ± 3.33%, η_p_^2^ = 0.45; Figure 3D, Experiment 4b: 1.96x10^5^ ± 3.46%, η_p_^2^ = 0.45) and there was some evidence that errors increased when distractors were attended for Experiment 3 only (Experiment 3: BF_10_= 10.81 ± 0%, η_p_^2^ = 0.21; Experiment 4a: 1.23 ± 3.37%, η_p_^2^ = 0.09; Experiment 4b: 1.22 ± 3.62%, η_p_^2^ = 0.07). Consistent with Experiments 1 and 2, evidence supports the null hypothesis that stimulus noise and distractors do not interact to increase memory errors (Experiment 3: BF_10_= 0.15 ±- 3.46%, η_p_^2^ = 0.007; Experiment 4a: 0.37 ± 3.61%, η_p_^2^ = 0.005; Experiment 4b: 0.35 ± 3.80%, η_p_^2^ = 0.007). A post hoc Bayesian t-test also suggests strong evidence that the presence of distractors increased WM errors compared to when there was no distractor (Experiment 3: BF_10_= 5.82x10^4^ ± 0%, Cohen’s d = 0.87).

**Figure 4.**
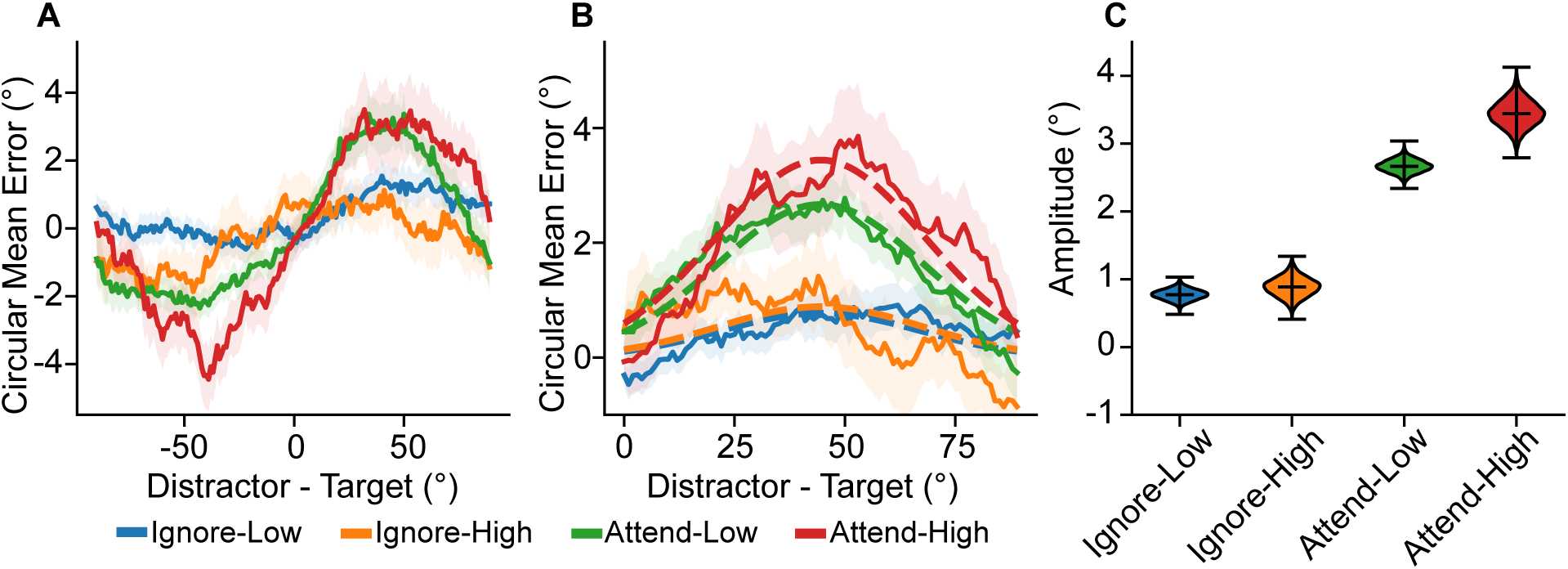
(A) Moving average distractor induced bias. Shaded region represents SEM. (B) Folded distractor induced bias (solid lines) with fitted Gaussians (dashed lines). (C) Amplitudes of Gaussian fits, center bar represents mean and out bars represent minimum and maximum.

#### Biases in memory error

The results shown in Figure 4 describe the Gaussian fits on circular mean error across conditions in Experiment 3 (4A-B) and the circular mean error across distractor-sample differences (4C) for Experiment 4. In Experiment 3, across all conditions, behavioral errors showed an attractive bias towards the distractor orientation. The magnitude of the bias was larger with higher stimulus noise (permutation test: *p* < .001) and when the distractor stimulus was attended (permutation test: *p* < .001). Finally, there was a marginally significant interaction between stimulus noise and distractor relevance (permutation test: p = 0.06).

As in Experiment 3, Experiment 4, which used four discrete target/distractor offsets as opposed to continuous sampling, also reveals systematic biases in memory errors. There is a non significant trend of a repulsive (as opposed to attractive) bias when the distractor is ignored (Experiment 4a: permutation test p = 0.51; Experiment 4b: permutation test p = .50) (Figure 5A-D). In contrast, attending to a distractor produced memory errors that were attracted to the distractor orientation (Experiment 4a: permutation test p < .001; Experiment 4b: permutation test p < .001). There is no significant effect of stimulus noise on bias (Experiment 4a: permutation test p=0.4; Experiment 4b: permutation test p = 0.36) and there was a significant interaction between noise and attention conditions in 4b only (Experiment 4a: permutation test p = 0.51; Experiment 4b: permutation test p = 0.01).

**Figure 5.**
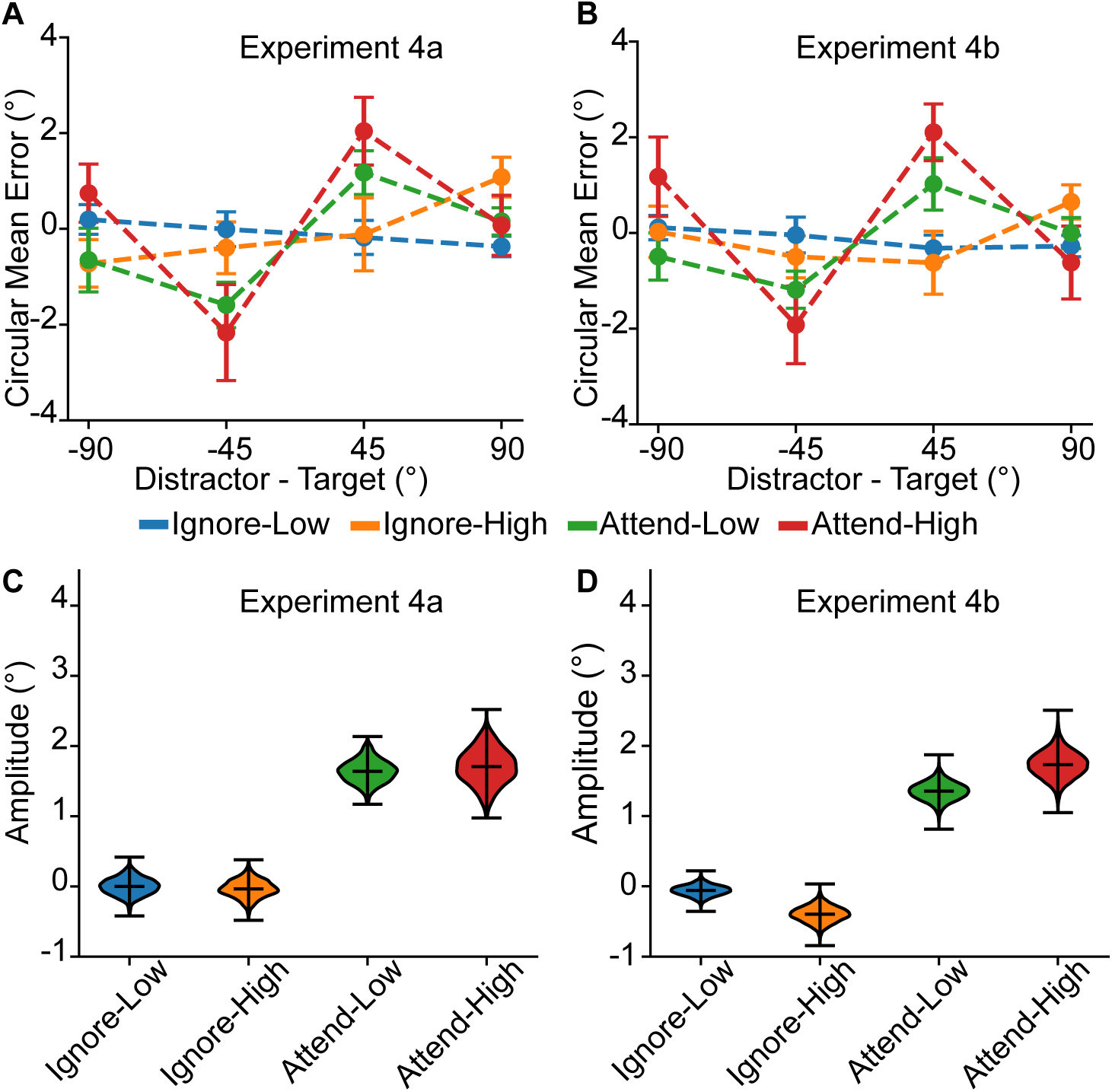
(A-B) Distractor induced bias. Error bars are SEM. (C-D) Estimated amplitudes for pseudo-subject samples, center bar is mean and outer bars are minimum and maximum.

## Discussion

When using memoranda and distractors that do not share a common feature, we find that stimulus noise and distractor presence and relevance exert largely additive influences on the overall magnitude of memory errors. However, using memoranda and distractors that share a common feature allowed us to more directly test for systematic interactions. We found that distractors bias patterns of memory recall error and that these biases are most pronounced when memory representations are noisy and when distractors are behaviorally relevant. While prior work has shown similar attractive biases (Dubé et al., 2014; Huang & Sekuler, 2010; Lorenc et al., 2018; Mallett et al., 2020; Nemes et al., 2011, 2012; Rademaker et al., 2015; Teng & Kravitz, 2019; Van der Stigchel et al., 2007; Zhang & Lewis-Peacock, 2023), it is unclear whether many of these effects are attributable to overlapping representations or whether the distractor influences post-perceptual stages of decision making (such as those proposed to support attractive serial dependence: Bliss et al., 2017; Papadimitriou et al., 2015; Sheehan & Serences, 2022; St John-Saaltink et al., 2016). In contrast, the systematic modulations reported here are consistent with a partially overlapping representational format for sensory and mnemonic information as manipulations that are expected to impact the quality of low-level visual information processing, such as more stimulus noise or attentional modulations of evoked neural responses to the distractor, lead to larger interactions between representations of seen and remembered stimuli. Consistent with these interactions playing out in early visual cortex, Hallenbeck et al. (2021) showed that both behavioral reports and neural representations in early visual areas V1–V3 were drawn towards nearby distractors (±12° from the remembered position) in a spatial working memory task.

Although reformatting of memory representations is a plausible encoding strategy to help reduce interference (Libby & Buschman, 2021; Xu, 2024), there is still little direct evidence that these non-interacting representations directly support memory recall. For example, Xu (2024) found that representations of remembered stimuli in parietal cortex did not change as a function of the identity of distractors presented during the delay period. In contrast, representations of remembered stimuli in early visual cortex did change as a function of distractor type, suggesting interacting sensory and mnemonic codes. Based on this observation, Xu proposed a privileged role for the untangled representations maintained in parietal cortex based primarily on a logical argument that these untangled representations should be less susceptible to interference and thus better able to support uncontaminated memory representations. Indeed, Xu (2024) did not observe any behavioral effects of distractor presence, a null result that is consistent with recall errors being based on untangled sensory and mnemonic representations. That said, Xu (2024) used a relatively small set of categorically distinct objects (e.g. coat hangers, bicycles, shoes) as stimuli, thus it is not clear what patterns of behavioral errors might arise or even whether to expect interference between such disparate stimulus classes. In contrast, we used a continuous feature space and a distractor that shared a common feature with the memory target, thus we could make precise predictions about how patterns of behavioral errors would change given interacting representations. The amplified attractive biases we observe with high stimulus noise and relevant distractors demonstrates that memory recall errors are not always based on representations that are recoded into abstract formats. Instead, in the present studies, behavior is biased towards incoming sensory inputs, consistent with at least partially overlapping codes for seen and remembered stimuli. That said, our stimuli are more artificial compared to the naturalistic objects used in Xu (2024), and using naturalistic stimuli can impact some facets of WM storage (Brady et al., 2016; Brady & Störmer, 2022; Chung et al., 2024; Morales-Torres et al., 2024; Thibeault et al., 2024). Future work might combine the advantages of well-parameterized artificial stimulus spaces and natural images by using tools such as CNNs to make predictions about how representations of real-world objects might interact at different levels of processing to assess the role of entangled (or untangled) representations on behavioral reports.

While our results indicate that behavioral errors can exhibit a bias consistent with interacting sensory and mnemonic representations, they do not imply that mnemonic representations are maintained in a “pure” sensory-like format. Instead, they establish some overlap in the representations that is more pronounced for weaker memory representations induced by sensory uncertainty and behaviorally relevant distractors (Figure 5A). In addition, changes in the priority of a memory or in the minimal amount of information about a complex stimulus that can efficiently support recall can lead to a recoding of sensory representations (e.g. collapsing a motion direction into a lower dimensional representation of orientation, or outsourcing a visual representation to motor cortex when the behavioral response is known in advance) (Duan & Curtis, 2024; Henderson et al., 2022; Kwak & Curtis, 2022; Yu et al., 2020). However, the present experiments require both high precision representations to support accurate recall and include feature- and behaviorally-relevant distractors. Therefore, relying on an abstract or lower-dimensional code is likely sub-optimal and a more sensory-like representation might better support performance, even at the cost of systematic biases. Taken together, recent work suggests that WM is supported by different types of representational formats as a function of the type of information being stored and concurrent task demands. Thus, instead of trying to identify a unified account of WM, these results further highlight the importance of considering how the mechanisms that support WM can flexibly adapt given changing task demands to strike a balance between providing adequate memory fidelity to achieve behavioral goals while using an efficient and compact code.

## Declarations

### Funding

This work was supported by the National Eye Institute award RO1-EY025872.

### Conflicts of interest

The authors have no competing interests to declare that are relevant to the content of this article.

### Ethics approval

All procedures were approved by the University of California, San Diego Institutional Review Board.

### Consent to participate

All human subjects provided informed consent.

### Consent for publication

Not applicable

### Author Contributions

Holly Kular: Conceptualization; Data curation; Formal analysis; Investigation; Methodology; Resources; Software; Validation; Visualization; Writing – original draft; Writing – review & editing.

John T. Serences: Conceptualization; Funding acquisition; Investigation; Methodology; Project administration; Resources; Supervision; Visualization; Writing – review & editing.

### Availability of data and materials & Code availability

No aspects of Experiment 1 and 2 were preregistered; all materials, data, and analysis scripts are available anonymously online (https://osf.io/xdte2/overview?view_only=32273d55e2c04ad787442758d0941c61). For Experiment 3 and 4, the design, sample size, exclusion criteria, and analysis plans were preregistered on *AsPredicted* (Experiment 3: https://osf.io/k76dh/overview?view_only=0fa0f580d7a443629c21c60cd780b763; Experiment 4: https://osf.io/eafkd/overview?view_only=c0b92b8534084773bae496c0fed21189) and all materials, data, and analysis scripts are available anonymously online (https://osf.io/xdte2/overview?view_only=32273d55e2c04ad787442758d0941c61).

## Notes

https://osf.io/xdte2/overview?view_only=32273d55e2c04ad787442758d0941c61

https://osf.io/eafkd/overview?view_only=291aecd88a6442028422e6c7018696f6

https://osf.io/k76dh/overview?view_only=081690d1f35d4140ac5ea4f1468efb14

### Competing Interest Statement

The authors have declared no competing interest.

